# Anisotropic Correction of Beam-induced Motion for Improved Single-particle Electron Cryo-microscopy

**DOI:** 10.1101/061960

**Authors:** Shawn Q. Zheng, Eugene Palovcak, Jean-Paul Armache, Yifan Cheng, David A. Agard

## Abstract

Correction of electron beam-induced sample motion is one of the major factors contributing to the recent resolution breakthroughs in cryo-electron microscopy. Improving the accuracy and efficiency of motion correction can lead to further resolution improvement. Based on observations that the electron beam induces doming of the thin vitreous ice layer, we developed an algorithm to correct anisotropic image motion at the single pixel level across the whole frame, suitable for both single particle and tomographic images. Iterative, patch-based motion detection is combined with spatial and temporal constraints and dose weighting. The multi-GPU accelerated program, MotionCor2, is sufficiently fast to keep up with automated data collection. The result is an exceptionally robust strategy that can work on a wide range of data sets, including those very close to focus or with very short integration times, obviating the need for particle polishing. Application significantly improves Thon ring quality and 3D reconstruction resolution.

---

In recent years single-particle cryo-electron microscopy (cryo-EM) has made groundbreaking advancements in obtaining atomic resolution structures of macromolecules that are either refractory to crystallization or difficult to be crystallized in specific functional states ^1–4^. Central to this success has been the broad application of direct electron detection cameras, especially the Gatan K2, which not only have significantly improved detective quantum efficiency (DQE) at all frequencies but also have a high output frame rate, typically 10 to 40 frames per second. Nowadays, cryo-EM images of frozen hydrated biological samples are recorded as dose-fractionated stacks of sub-frames (movies)^5–7^, which enable correction of beam-induced sample motion that blurs the captured images^6,8^. Combining high sensitivity imaging, motion correction and advanced image processing and 3D classification strategies have made it practical to routinely determine three-dimensional (3D) reconstructions at near atomic resolution by single particle cryo-EM even for the most challenging samples^9–11^.

Sample illumination with the high-energy electron beam breaks bonds, releases radiolysis products and builds up charge within the thin frozen hydrated biological samples during image recording. The result is a combination of physical and optical distortions that can significantly deteriorate sample high-resolution information through image blurring. This has been one of the major factors limiting the achievable resolution of single particle cryo-EM^12^. The concept of recording the image as movie to correct sample motion was proposed long ago^13,14^, but only became practical after direct electron detection cameras were made available^7^. The fast sample motions can be measured by tracking the movement of images captured in a series of snapshots, either as the whole frame or as individual particles. Image motion can then be corrected by registering identical features in the sub-frames to each other, followed by summing the registered sub-frames to produce a motion-corrected image^6^. From the earliest studies it has been clear that the apparent sample motions can be anisotropic and idiosyncratic, such that the pattern of motion varies across the image. While in principle tracking should be straightforward, the practical challenge is the extremely low signal-to-noise ratio (SNR) in each individual sub-frame. Except for very largest particles, accurate motion measurement generally requires correlating the motion between sub-frames over large areas. That said, even sub-optimal motion correction can significantly restore high-resolution signals and improve the resolution of final 3D reconstructions^6,8^.

We previously developed an algorithm that made use of redundant measurements of image shifts between all sub-frames to derive a least squares estimate of relative motions between neighboring sub-frames. This algorithm, implemented in the program MotionCorr, provided an efficient correction of image motions with sufficient accuracy^6^ to enable the determination of numerous near atomic resolution 3D reconstructions^11,15^. Around the same time or soon afterwards, a number of different strategies were devised that either assume particles located nearby have similar motions or assume uniform motion of the entire frame or patches of the frame. Programs based on the former assumption include RELION that provides a movie-processing mode but uses a 3D reconstruction to track particle motions^8,16^, Xmipp that implemented an Optic Flow algorithm^17^, and *alignparts*_*lmbfgs* that implemented a regularized Fourier Space optimization algorithm to track neighboring particles^18^. Programs based on tracking the full frame or parts of the frame include MotionCorr^6^ and the iterative whole frame alignment procedure used in electron tomography image alignment^19^. All of these algorithms have demonstrated the ability to recover high-resolution signals to varying degrees and have improved the resolutions of 3D reconstructions.

Ideally, single particle cryo-EM images should be acquired with the smallest possible defocus to enhance high-resolution information, and with the shortest sub-frame exposure times to reduce motion trapped within individual sub-frames, in particular for the first few frames where the sample has the least radiation damage but moves most rapidly^20^. Additionally, motion detection should be done on the smallest possible local area to best capture the anisotropic motion. Unfortunately, each of these constrains would significantly reduce the SNR in each subframe, ultimately leading to incorrect estimates of the beam-induced motion. For example, previous experiments with MotionCorr revealed that subdividing the images into areas smaller than ~2000 × ~2000 pixels or going to sub-frame integration times of less than 100 milliseconds worsened resolution due to increased errors in motion tracking.

Another recent advance, where a model of radiation damage is used to weight the individual sub-frames in Fourier space, allows collection to very high dose (80-100e^−^/Å^2^) producing a high contrast image without degradation of the high-resolution signal^21^. While having a single correctly weighted high contrast image has clear advantages for particle picking, orientation refinement, and overall computability, taking advantage of this strategy requires that each of the sub-frames be optimally aligned over the full image area. Therefore, improving motion correction algorithms is practically very important for advancing single particle cryo-EM such that the highest resolution information is extracted from each image.

In this work we experimentally validated the doming behavior that was proposed to describe beam-induced sample motion^5^ and discovered that the initial motion begins at surprisingly low doses. Our observations supported a model in which the sample is smoothly and continuously deformed throughout the exposure. The extreme sensitivity to dose emphasizes the need to acquire at very short sub-frame durations, further raising the demands for performance at low SNRs.

To meet the compounding demands for locality and low SNR, it was not possible to accurately determine local shifts by comparing noisy sub regions of each frame to one another as done within MotionCorr. Instead, we compare local image regions to an approximated sum for each image subregion, and then iterate translation determination to improve the sum. The resultant shifts are further constrained both spatially and temporally as informed by the physical behavior of the sample. This is accomplished by deriving the final shifts from a fit of a time-varying two-dimensional (2D) polynomial function to the local motions derived from the different patches of the image. Each image sub-frame is subsequently remapped using this smooth distribution of beam induced motions at each individual pixel and summed with or without radiation damage weighting. This algorithm has been implemented as MotionCor2, a parallel computing program running on Linux platform equipped with multiple GPUs. Our tests have shown that this program is very robust, sufficiently fast to keep up with automated data collection and capable of significantly extending the resolutions obtainable by single particle cryo-EM.

## RESULTS

### Modeling of beam-induced motion

Early studies suggested that beam induced motion can be described as specimen doming^5^. This model predicts a significant sample motion perpendicular to the sample plane (z-motion, Figure 1a), which, when the sample is tilted, should be projected onto the image plane as a motion perpendicular to the tilt axis (Figure 1b). We explored this by directly measuring motions within movie stacks recorded from specimens tilted at various angles. We collected a number of dose fractionated tomographic tilt series from various frozen hydrated biological samples. By using rather low magnifications (4.3Å/pixel), we could accurately determine full frame shifts with doses down to 0.05 e^−^/Å^2^ using MotionCorr^6^. Representative traces of uniform whole frame motion of the same specimen at three tilt angles α=−30°, 0°, and +60° clearly show that the measured motion is much larger at higher tilt angles and mainly perpendicular to the tilt axis (Figure 1c), supporting the dome model. Consistent with the earlier study ^5^, the motion within the very first 0.2 e^−^/A^2^ is substantial. The beam-induced shifts observed during normal single particle cryo-EM must be a combination of sample drift, residual tilt, and non-idealities in the doming such that local motions are not purely along Z.

**Figure 1.**
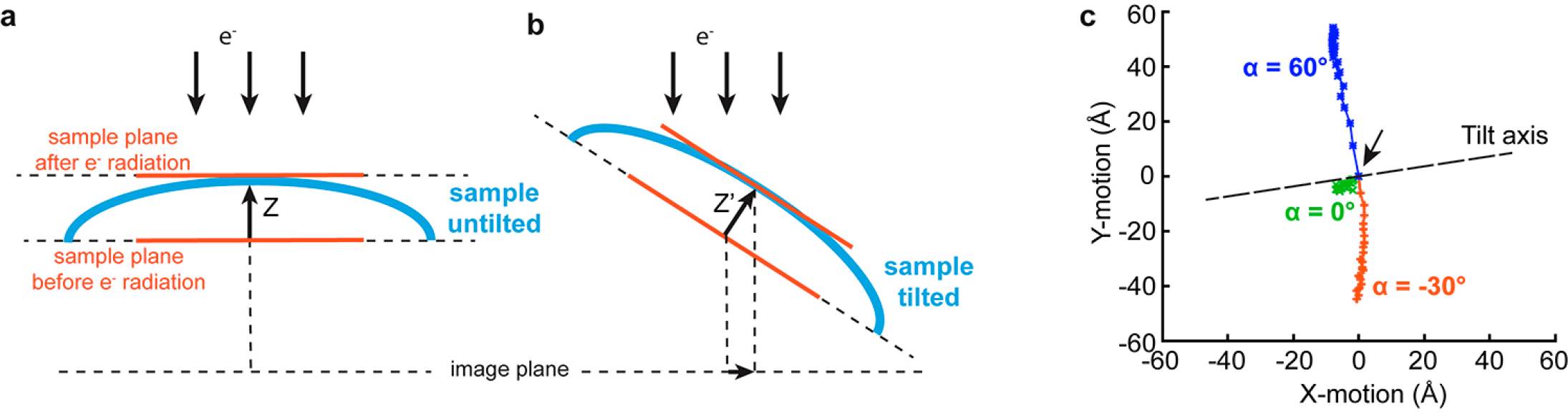
*Doming model that describes the motion of frozen hydrated samples induced by illuminating high-energy electron beam*. (a) Schematic drawing illustrates the doming of the sample without tilting under electron beam. The major motion (z-motion) is parallel to the incident beam and thus its projection to the image plan produces little in-plane shift. (b) When the sample is tilted, projections of z-motion produce observable motions in the image plan. (c) Traces of the projected motion measured at three different tilt angles (α-angle) extracted from a dose fractionated tilt series acquired on a frozen hydrated specimen of a wild-type Drosophila melanogaster γ-Tubulin Ring Complex. The black arrow in the plot indicates the starting position of the motion.

### Motion correction using polynomial constraints

Whole frame based motion correction algorithms, such as ^6,19^, correct only the uniform motion across the entire image frame, leaving the residual non-uniform local motion uncorrected. A common strategy for correcting local motions is to spatially partition a movie stack into multiple stacks of patches where the correction is performed individually^6^. Ideally, the smaller the patch size is, the better locality of motion correction. In practice, there is a limit on patch size, below which the motion cannot be corrected with sufficient accuracy due to insufficient SNR. Furthermore, dividing the whole image into a limited number of patches and correcting motion within each patch independently causes edge artifacts at the boundaries of patches in the corrected image, the so-called checkerboard artifact. This not only creates additional challenges in single particle work, but obviates utilization for tomography.

To avoid these problems, we chose to combine iterative patch-based motion measurement with restraints derived from the physical behavior to obtain a function that describes the motion of each pixel across the entire image. Since a dome can be geometrically approximated by a quadratic surface, we chose to fit the locally measured shifts from all patches to a time-varying polynomial function that is quadratic in *xy* plane and cubic with respect to exposure time *t*.

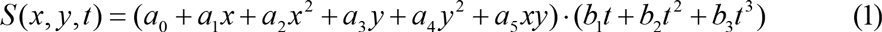

where, *S* is the shift at the integer pixel coordinate *x* and *y*. Since this function is second-order differentiable, a smooth estimate of the shift can be obtained at each pixel of each sub-frame. Not only does this strategy eliminate checkerboard artifacts, but importantly it also provides a more thorough correction of image motion. It effectively dampens the fluctuations of local measurements influenced by image noise.

### Bad Pixel correction

Despite the considerable efforts of the camera manufacturers, there are often image defects that remain uncorrected by their calibration methods. In the case of our Gatan K2 cameras, we observe a variety of imperfections including hot pixels, cold pixels and clusters of 6 × 6 superresolution pixels having distorted intensities (Supplementary Fig 1a). As the intensity distribution is similar to a 2D truncated sinc function (or an inverted version), these are likely the consequence of a single pixel defect perturbed by Gatan’s electron counting/super-resolution algorithm. It was also observed that the locations of the defects vary from exposure to exposure (Supplementary Fig 1b and c). The longer the exposure, the more clusters are present. To minimize possible deleterious effects of these systematic image errors on the low SNR image registration, we first detect these defects based on the image statistics within each stack. Defects are identified first from the sum of gain-corrected sub-frames without motion correction, with the clusters located by cross-correlating a template 2D truncated sinc function. Once identified, these defects are corrected for each sub-frame using values of good pixels randomly selected from the neighborhood. Only the defect-corrected stacks are used for subsequent motion correction.

### Motion correction program MotionCor2

We first correct the uniform global motion by iteratively aligning each sub-frame against a reference that is the motion corrected sum of all other sub-frames based upon the translational alignment obtained in the previous cycle. Excluding the sub-frame being aligned from the reference prevents a strong auto-correlation peak at the origin that may influence the determination of the real cross correlation peak. The measured sub-frame shifts at each iteration are the residual errors from the last iteration, and the alignment procedure is terminated when the maximum residual error is below a specified tolerance. The measured global shifts are then corrected by phase shifting in the Fourier domain to yield a global-motion-corrected stack. The image stack is then divided into a grid of patches, and the same iterative alignment procedure is performed on each patch. This provides the local motion at a series of discrete locations for each time point within the exposure. These patch based local motions are then fit to polynomial functions defined in (1), *S_x_*(*m,n,t*) for *x* shifts and *S_y_*(*m,n,t*) for *y* shifts. As a result, *x* and *y* shifts can be calculated according to the fit functions at each pixel (*m*, *n*) in each sub-frame. Since the calculated shifts at individual pixels are typically non-integer, the corresponding correction requires interpolation in the real space. To minimize the attenuation of high-resolution signals due to interpolation, bilinear interpolation is performed on super-resolution pixels. The final image is obtained by cropping the corrected sum in the Fourier domain to the user specified resolution. In practice the whole stack is typically divided into 5×5 partitions, an empirical choice that provides a good balance between precision and efficiency.

### Integration of dose weighting

Dose weighting each motion-corrected sub-frame according to expectations from analysis of radiation damage allows cryo-EM images to be recorded with significantly higher total electron doses. This improves low-resolution image contrast without sacrificing high-resolution SNR^21^. We have integrated this weighting scheme into MotionCor2 to streamline all the necessary preprocessing steps prior to the normal cryo-EM processing pipeline.

For biological samples embedded in vitreous ice, radiation damage dampens high-resolution signals that are useful for high-resolution structure determination, hence the need for the weighting. While the dose-weighting scheme appropriately down-weights the high-resolution biological information from the high dose data within the summed image stack, it also unnecessarily attenuates the high-resolution Thon ring signals used for CTF determination. To avoid this undesired side effect, MotionCor2 generates both a dose-weighted summed image for cryo-EM reconstruction and an un-weighted summed image for CTF estimation (Supplementary Figure 2). In order to maximize the throughput of motion correction, parallel computation was implemented in three levels in MotionCor2 on a Linux platform equipped with multiple GPUs.

### MotionCor2 improves accuracy of motion correction

We tested the performance of MotionCor2 using two previously acquired single particle cryo-EM datasets: the archaeal 20S proteasome and the rat TRPV1 ion channel. Both were previously processed using whole frame based MotionCorr and produced near atomic resolution structures^6,11^. These two datasets were collected using the same microscope settings and similar electron dose rates on the camera. As these were older data sets the total dose was limited: the 20S stacks contained 25 sub-frames (total dose of ~34 electrons per Å^2^) whereas the TRPV1 stacks had 30 sub-frames (total dose of ~41 electrons per A^2^). We re-processed motion correction for these two datasets using MotionCor2, configured to run a 5×5 patch.

The trajectory of the full frame (global) motion determined by the original MotionCor program is shown in Figure 2a. Because of the very high SNR for these images, a very similar global motion trace was determined using the MotionCor2 iterative strategy (long trace, Figure 2b). The measured local motions for each patch (traces in each panel, Figure 2b) in this representative image stack are qualitatively similar to the smoothed trajectories once the physical constraints are applied (Figure 2c) indicating that the added robustness does not come at the cost of capturing local motion. The new algorithm improves Thon ring signals at high-resolution and leads to a better correlation with the simulated contrast transfer function (CTF) (Supplementary Figure 2a). Over the entire data set, almost all images showed a significant improvement in recovery of high-resolution Thon ring signals (points below the dashed line, Figure 2d and Supplementary Figures 3a, b and c).

**Figure 2.**
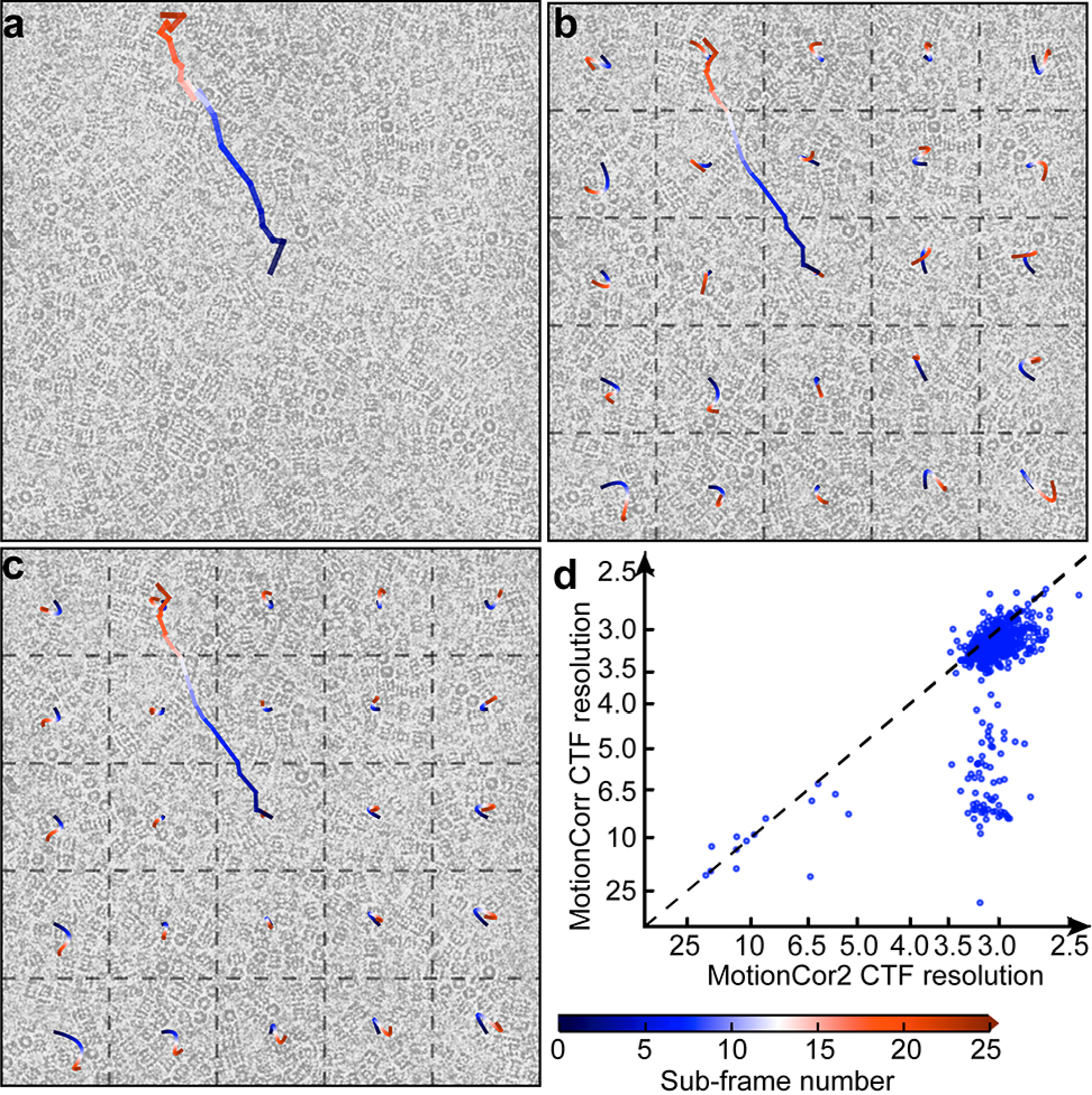
*Motion correction of cryo-EM images by MotionCor2*. (a) Image of frozen hydrated archaeal 20S proteasome overlaid with the traces of motions determined. The long trace originated from the center of image is the global motion based upon full-frame alignment. The whole frame is divided into 5 × 5 patches, and traces of each patch are predicted from the polynomial function. (b) The same image overlaid with the motion traces of each patch as determined from MotionCor2. (c) The trace of globe motion determined from the same image using the original MotionCorr. (d) Resolutions of micrographs estimated using CTFFIND4 ^22^ from the correction by both MotionCorr (vertical axis) and MotionCor2 (horizontal axis). Circles on the lower right side of the dashed line represent micrographs which MotionCor2 produced better correction.

Using maximum-likelihood based refinement and reconstruction algorithms, we redetermined the 3D reconstruction of the archaeal 20S proteasome from images that were processed both by the whole frame motion correction strategy originally used (MotionCorr) and the algorithm presented here (MotionCor2). Using MotionCor2, the nominal resolution of the 3D reconstruction of archaeal 20S proteasome was improved from 2.73A to 2.58Å (Figure 3). The visible structural features were also significantly improved (Fig. 3 and Supplementary Figure 3d and e). Most backbone carbonyls are now clearly visible, as well as the precise rotameric states of amino acid side chains (Figure 3b and c). Such improved structural detail would clearly facilitate more accurate model building. Indeed, after real-space refinement of our previous atomic model into the new map, we have substantially improved model validation statistics such as the EMRinger score, the real-space cross correlation, and various model geometry scores given by MolProbity (Supplementary Table 1).

**Figure 3.**
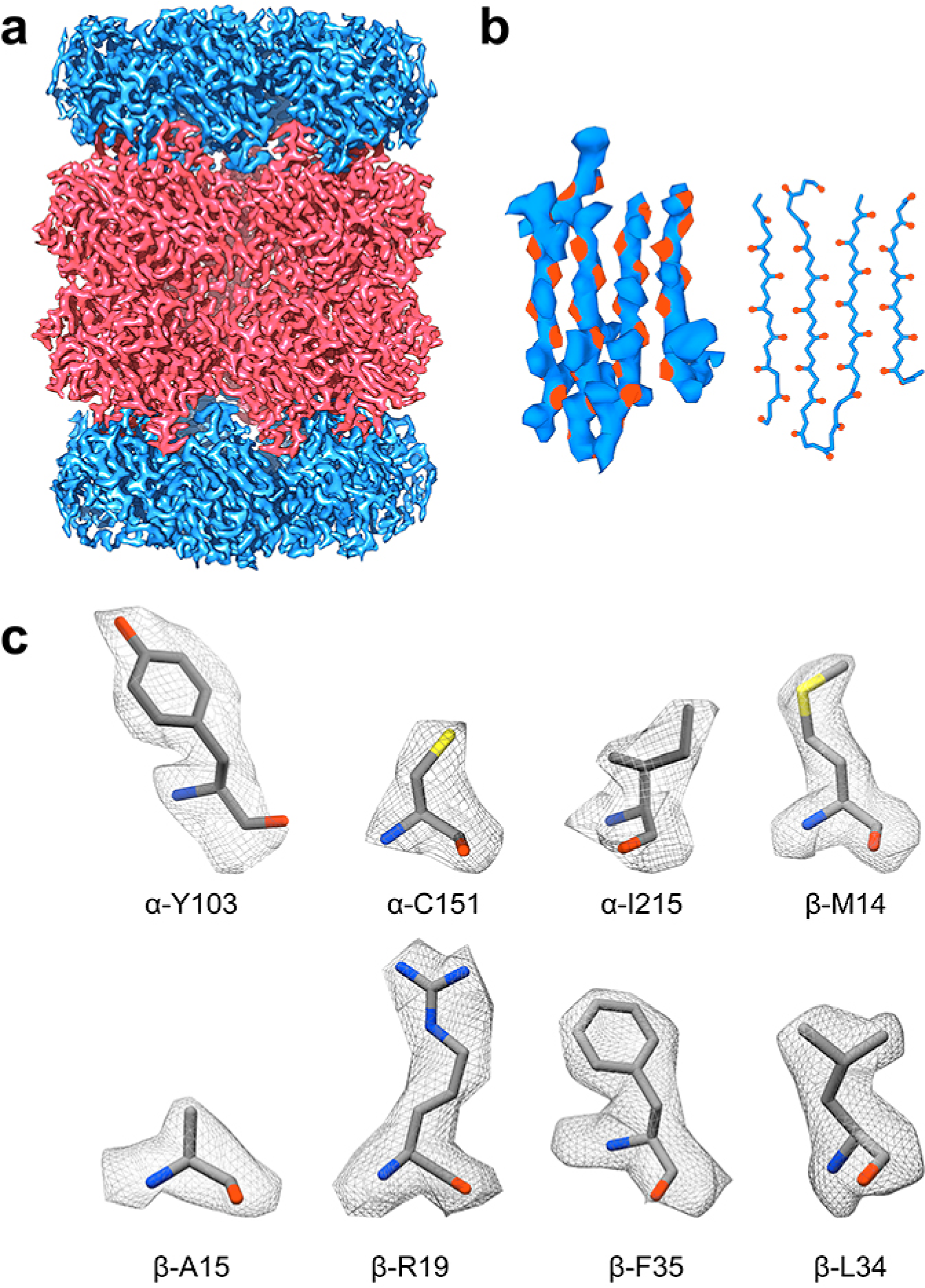
*3D reconstruction of archaeal 20S proteasome at 2.5Å resolution*. (a) 3D reconstruction of archaeal 20S proteasome filtered to 2.58Å resolution and sharpened by a temperature factor of −103.8Å^2^. (b) Density of a β-sheet, where the main chain carbonyl is colored in red. Visualization of main chain carbonyl density requires better a resolution that is better than 3A. (c) Representative side chain densities.

Similarly, we re-processed our previous raw micrographs from the TRPV1 ion channel ^11^ and re-determined its 3D reconstruction. MotionCor2 improves the nominal resolution from 3.5Å to 3.15Å (Supplementary Figure 4). Such improvements are particularly obvious in some trans-membrane regions, where extra densities associated with the TRPV1 protein are now seen to have well-defined features that can only now be interpreted as lipid molecules (Supplementary Figure 4e and f). Thus, for both 20S proteasome and TRPV1 datasets, MotionCor2 produced noticeable resolution improvements.

We also tested the performance of MotionCor2 with a new dataset of archaeal 20S proteasomes collected from our TF20 microscope operated at 200kV equipped with a Gatan K2 camera. The sub-frame exposure time was set to 75ms, approximately one third of the frame exposure time (200ms) we typically use. About half of these images were recorded with defocus set between −0.4μm to −1.0μm. A representative image collected with −0.5μm defocus is shown in Figure 4 and MotionCor2 restored its Thon rings to close to 3Å resolution. We picked a small set of 36,000 particles from motion corrected images and determined a 3.4Å reconstruction of the 20S proteasome (Supplementary Figure 5). Interestingly, a reconstruction of 4.1Å resolution could be determined using only the 3,297 particles having defocus values less than −1 μm (Supplementary Figure 5a and d). In contrast, using the same number of particles, but higher defocus (−2 ~ −2.5 μm), the resolution of the 3D reconstruction is 4.9Å (Supplementary Figure 5a and e). This emphasizes the importance of being able to accurately motion correct very low defocus images.

**Figure 4.**
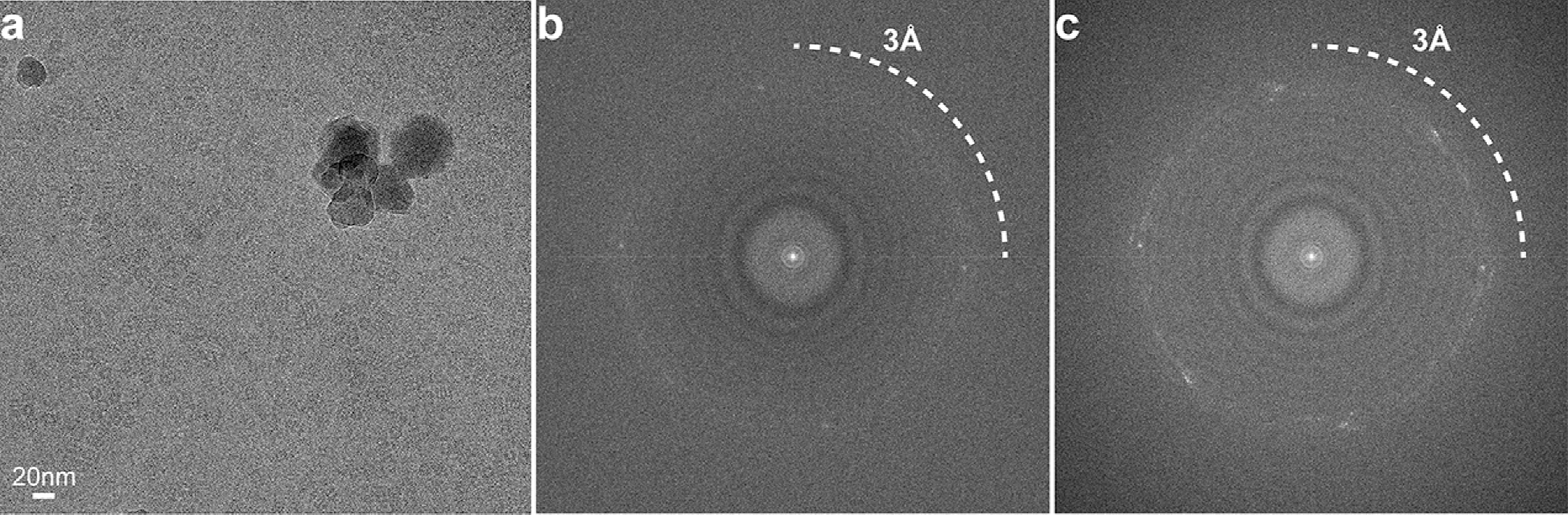
*Motion correction of low defocused image*. (a) A micrograph of frozen hydrated archaeal 20S proteasome recorded with a defocus of −0.4 μm. (b) and (c) Fourier power spectra are shown from the same image before motion correction (b), and after motion correction with MotionCor2 (c). The motion correction of the same micrograph using MotionCorr did not produce satisfactory result.

## DISCUSSION

Improving both accuracy and efficiency of motion correction has been an extensive topic of research since the direct electron detection camera and dose-fractionated imaging techniques were first introduced to cryo-EM ^6,8,16–19^. Compared with algorithms that only correct the global motion, the new algorithm implemented in MotionCor2 has shown consistant and significant improvements while maintaining superb robustness. Because image motions are corrected at the pixel level prior to any further image processing, it is practical to take advantage of the benfits of dose-weighting without the computationally intensive steps that track and correct motion of individual particles implemented in RELION ^16^. Importantly, our full frame motion correction strategy is also applicable to tilted data collected by cryo tomography. Furthermore, by outputting both unweighted and dose-weighted frame averages, CTF determination is optimized. Particles boxed from the weighted images have enhanced low-resolution contrast and high-resolution SNR, facilitating more accurate particle alignment and classification, thereby producing 3D reconsturctions having improved resolutions.

Our tests have shown that the new algorithm also works well with dose-fractionated image stacks recorded with defocus and frame exposure times that are significantly lower and shorter than commonly used (Figure 4). This could potentially impact the collection of single particle cryo-EM datasets when aiming for the highest resolution possible. For atomic-resolution single particle cryo-EM reconstruction, there are many benefits of recording cryo-EM images with low defocus, preferrable less than −1 um ^20^. Provided that image motion can be corrected accurately, dose weighting allows images to be recorded with sufficently high total dose to enable processing while realizing the benefits of low defocus to obtain high resolution. Another practical issue is that motion is most rapid and most anisotropic in the first few sub-frames, resulting in substantial deterioration of high-resolution information within the first ~ 3 e^−^/Å^2^ of dose. While these have typically been excluded or substantially down weighted, we have shown that recording images with shorter sub-frame exposure times can reduce such deterioriation, again, provided that the local image motion captured in the short sub-frames can be corrected (Supplementary Figure 5b). Our tests have shown that MotionCor2 provides satisfactory corrections to images recorded with both low defocus and short frame exposure times (Figure 4). Such robust motion correction enabled determination of a 4Å resolution reconstruction using only ~3,000 particles from low-defocused images with short frame exposure times recorded on an F20 microscope. This dramatically enhances the utility of such an instrument for routine single particle cryoEM.

## Acknowledgements

We thank X. Li for helpful discussion during the initiate stage of this work. We also thank M. Braunfeld for supporting the cryo-EM facility at UCSF, and C. Kennedy for supporting the computational infrastructure for processing cryo-EM data. This work was supported in part by grants from National Institute of Health, GM031627 to D.A. and P01GM111126, P50GM082250, R01GM082893 and R01GM098672 to Y.C. Y.C. and D.A. are Investigators of Horward Hughes Medical Institute.

---

**Supplementry Figure 1.**
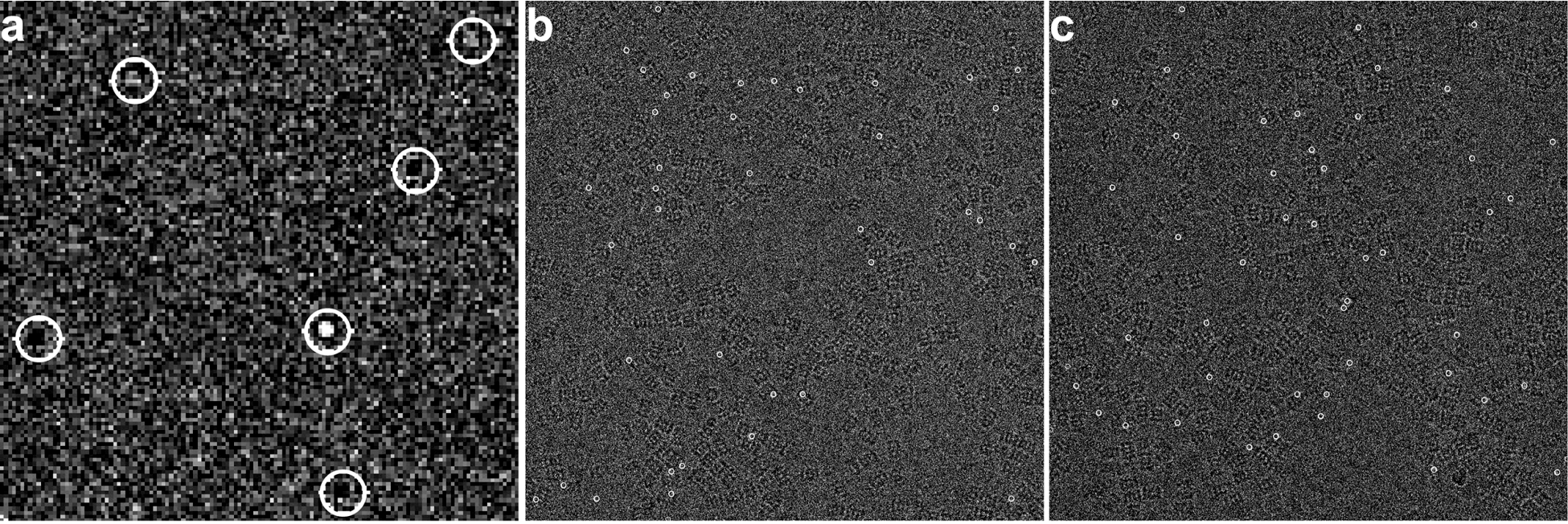
*Defect pixel fixation in MotionCor2*. (a) An example showing defected pixels (marked with white circles) in image captured with K2 Summit camera. (b) and (c) Two different images collected one after the other show defect pixels (marked with white circles) in different locations.

**Supplementary Figure 2.**
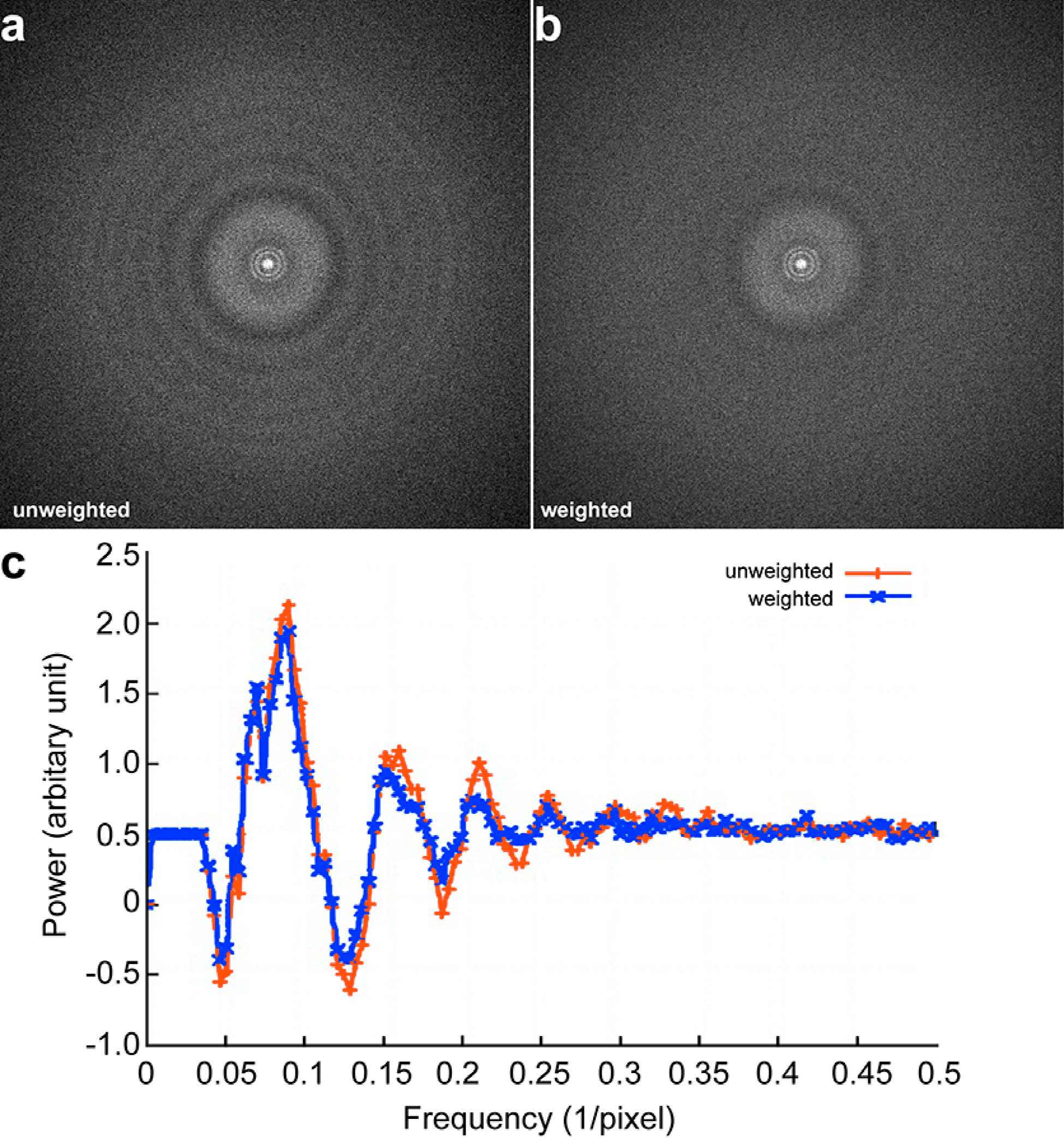
*Influence of dose weighting on Fourier power spectrum*. (a) and (b) Fourier power transform calculated from dose-weighted (a) and un-weighted (b) image after motion correction. (c) The rotation averages of dose-weighted (blue) and unweighted (red) Fourier power spectra shown in (a) and (b).

**Supplementary Figure 3.**
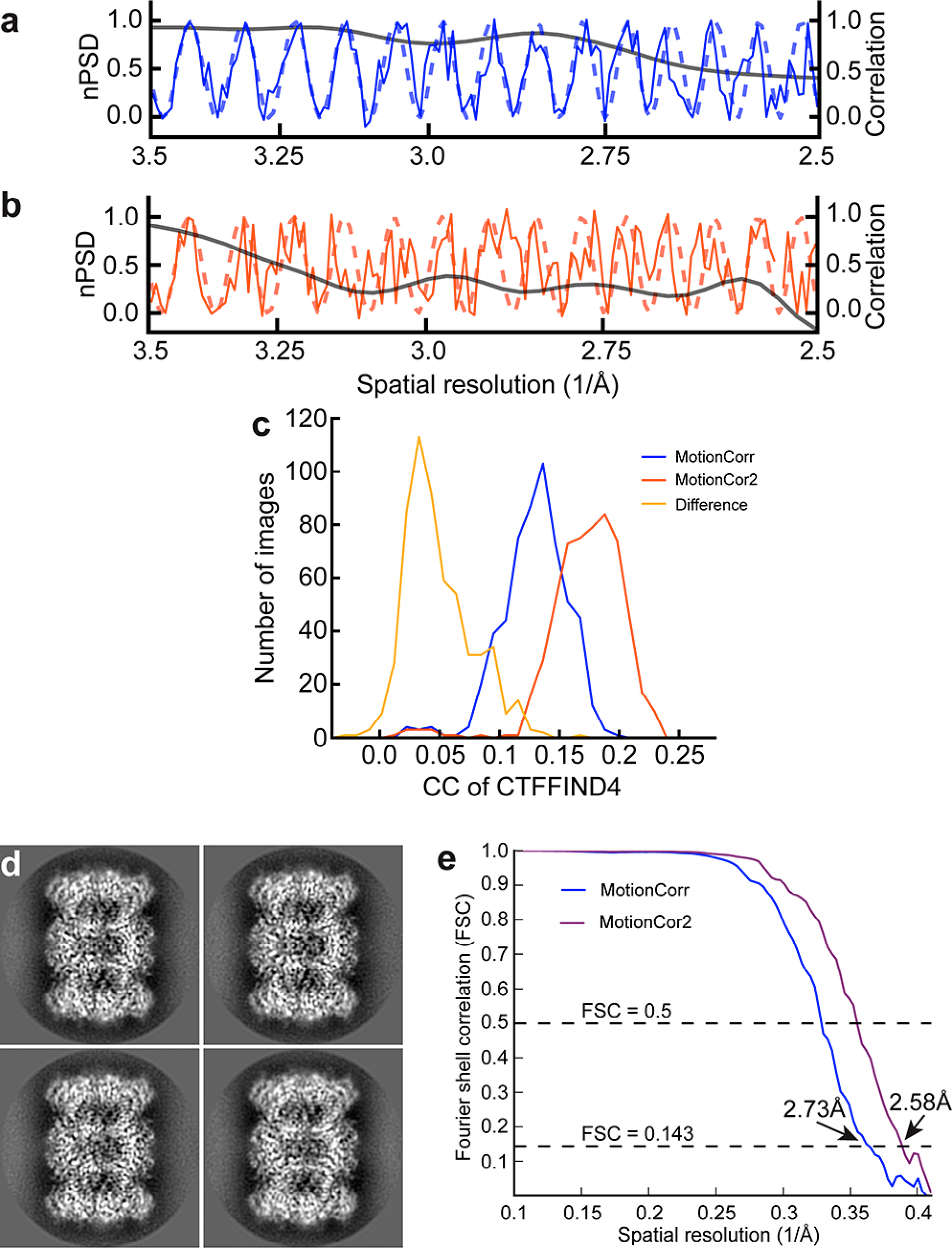
*Comparison of motion corrections by MotionCor2 and MotionCorr on 3D reconstruction of archaeal 20S proteasome*. (a) Rotationally averaged Fourier power spectrum of image after motion correction by MotionCor2 (dashed line) and fitted contrast transfer function (solid line). (b) Rotationally averaged Fourier power spectrum of image after motion correction by MotionCorr (dashed line) and fitted CTF (solid line). Solid black line in both (a) and (b) indicate cross correlation coefficient between the rotationally averaged Fourier power spectrum of image and fitted CTF. (c) Histogram of cross correlation coefficients between calculated and simulated Fourier power spectrum of MotionCorr corrected image (blue) and MotionCor2 corrected image (red). The difference, which shows the amount of improvement, is shown in yellow. (d) Selected 2D class averages of 20S proteasome. (e) Fourier Shell Correlation (FSC) curves of 3D reconstruction determined using micrographs corrected by MotionCorr (blue) and MotionCor2 (purple).

**Supplementary Figure 4.**
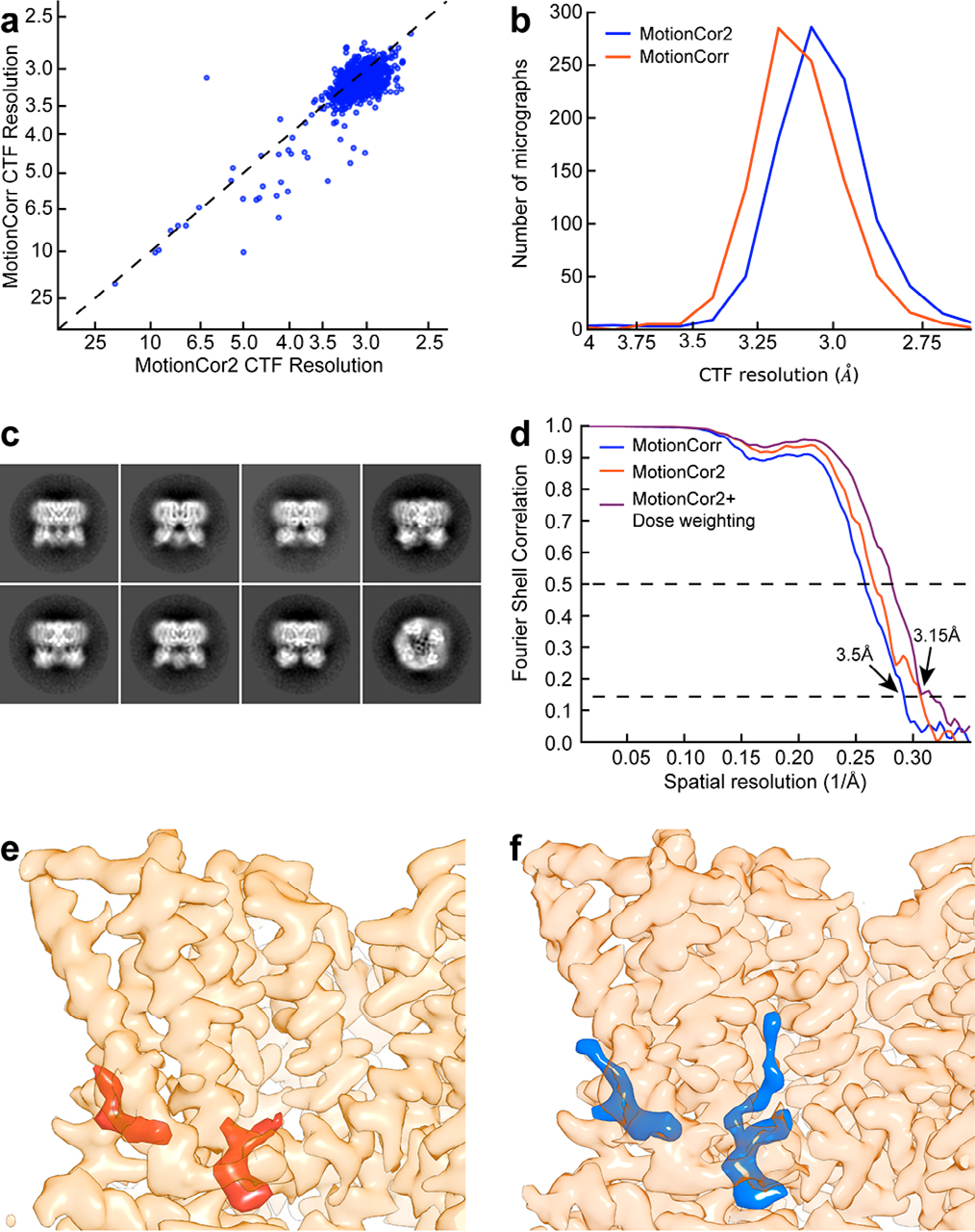
*Comparison of motion correction by MotionCor2 and MotionCorr on 3D reconstruction of rat TRPV1 ion channel*. A published dataset of frozen hydrated rat TRPV1 ion channel was reprocessed using MotionCor2. (a) Resolutions of micrographs estimated using CTFFIND4 ^22^ from the correction by both MotionCorr (vertical axis) and MotionCor2 (horizontal axis). (b) Histogram of cross correlation coefficients determined by using CTFFIND4 using image corrected by MotionCorr (blue) and MotionCor2 (red). The difference, which shows the amount of improvement, is shown in yellow. (c) Representative 2D class averages of frozen hydrated TRV1 particles. (d) FSC curves of 3D reconstructions determined from the same data set after motion correction by MotionCorr (blue) and MotionCor2 (red). (e) A representative view of the TRPV1 ion channel generated from previously published density map ^11^. (f) The view of the same region of TRPV1 density map determined after re-process motion correction using MotionCor2.

**Supplementary Figure 5.**
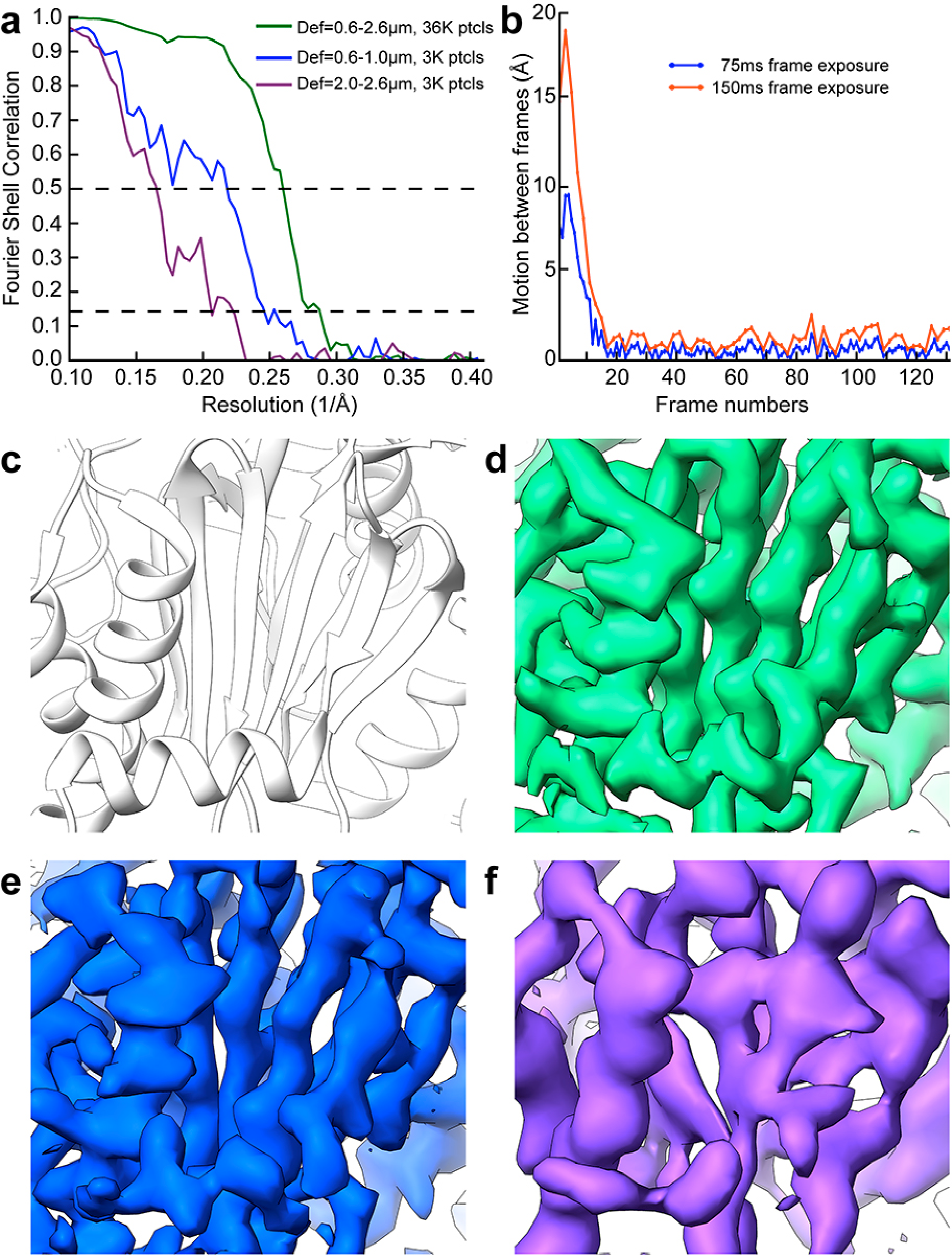
*3D reconstruction of archaeal 20S proteasome from 200kV electron microscope*. (a) Red: FSC curve of an archaeal 20S proteasome 3D reconstruction determined from a dataset of ~36,000 particles collected with a TF20 electron microscope operated at 200kV acceleration voltage. The defocus range was set between 0.6μm and 2.6μm. Blue: FSC curve of a 3D reconstruction using a subset of 3,000 particles with low defocus (0.6μm to 1μm). Purple: FSC curve of a 3D reconstruction of using another subset of 3,000 particles high defocus (2.0μm to 2.6μm). (b) A plot of motion between neighboring frames when frame exposure was set to 0.075 second (blue) and 0.15 second (red, by average two adjacent frames). (c) Ribbon diagram of a part of archaeal 20S proteasome. (d) Same region of the 3D reconstruction determined from the entire dataset, corresponding to red FSC curve in (a). (e) Same region of the 3D reconstruction determined from the subset of 3,000 particles with only low defocused particles (blue FSC curve). (f) Same region of the 3D reconstruction determined from a subset of 3,000 particles with only high defocused particles (purple FSC curve).

**Supplementry Table:** Comparison of atomic models of archaeal 20S proteasome (T. *acidophilum)* refined against the 3D density maps determined previously^6^ and in this work.

|  | Previous work | Current work |
| --- | --- | --- |
| Total number of particles: | 129,008 | 187,011 |
| Map resolution (FSC=0.143) | 3.3Å | 2.58Å |
| Molprobity score | 2.55 | 1.81 |
| Ramachandran outliers | 1.9% | 0.00% |
| Ramachandran favored | 90.10% | 96.2% |
| Rotamer outliers | 9.30% | 0.6% |
| C-beta outliers | 574 | 0 |
| Clash-score | 5.11 | 10.79 |
| Real-space cross correlation | 0.75 | 0.76 |
| EMRinger score | 3.08 | 3.95 |

## ONLINE METHODS

***Motion correction algorithm*** Image motion can be decomposed into two components, the uniform global motion and the non-uniform local motion that is described as the projection of the doming motion. The motion correction is divided into two steps, globe motion correction first followed by local motion correction.

The global motion is corrected iteratively by calculating cross correlation against the sum of all other sub-frames based upon the alignment of previous iteration. To avoid self-correlation, the sub-frame being measured is excluded from the sum. The measured shifts, i.e., the residual errors of the previous iteration of alignment, are corrected by shifting the phase in the Fourier transforms of sub-frames. The iteration stops when the residual errors are below a specified tolerance (typically, 0.5 super-resolution pixel) or a specified maximum number of iterations have reached.

The global-motion-corrected stack is partitioned into non-overlapping patches. The same alignment procedure is applied to each stack of patches. The local motion is therefore measured at a series of discrete spatial locations represented by the centers of patches. The measured local shifts of each patch of each sub-frame are fit to a polynomial function given in Eq. (1) that is quadratic in *xy* plane and cubic with respect to time *t*], which is the time stamp of a sub-frame during the exposure.

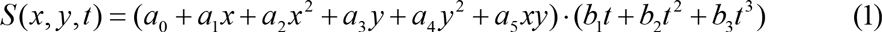

This equation can be expanded into Eq. (2) as follows.

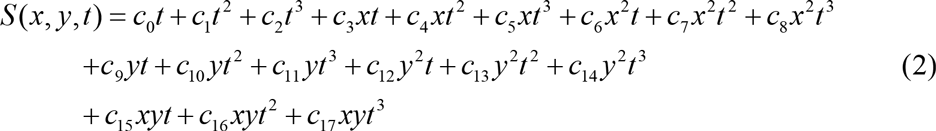

Least square fitting is used to determine the unknown coefficients *c_i_*. The measured *x* and *y* shifts are fit separately to Eq. (2), resulting two polynomial functions, *S_x_*(*x,y,t*) and *S_y_*(*x,y,t*), corresponding respectively to the *x* and *y* components of local motion. The vector function (*S_x_, S_y_*) provides a complete and smooth description of the local motion field, allowing smooth correction of the local motion at pixel level without inflicting edge effect. The curl of the vector (*S_x_, S_y_*) given in Eq. (3) is a time-varying linear function in (*x*, *y*) domain after applying partial derivatives to Eq. (2). Therefore, the fitted vector function (*S_x_*, *S_y_*) can also describe image rotation as observed by ^5^.

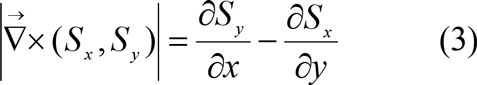

Since the calculated shifts are decimal and varying from pixel to pixel, the correction of pixel shifts are performed in real space using bilinear interpolation. To minimize the attenuation of high-resolution signals due to interpolation, the interpolation is performed on super-resolution pixels. The final image is obtained by cropping the corrected sum in Fourier domain to the user specified resolution.

**Parallel implementation.** The dome-model based motion correction algorithm was implemented in a GPU (Graphics Processing Unit) accelerated program named MotionCor2 that runs on a Linux platform equipped with one or more GPUs. High-throughput motion correction was achieved by means of parallel computation implemented in three levels. At the lowest level, pixel-wise computations, such as correction of gain reference to each pixel of a sub-frame, are implemented in various CUDA kernels. At the second level, independent operations such as Fourier transform of sub-frames and alignment of individual patches are distributed to all participating GPUs. The last level of parallelization is the batch processing of multiple movie stacks. The disk IO operations, i.e. loading a new stack and saving a corrected image, are performed in parallel with the intensive computation involved in correcting a loaded stack. This strategy minimizes the idle time of GPUs in waiting for a new stack. We assessed the efficiency of MotionCor2 configured for 5×5 patch-based motion correction on a Linux platform equipped with 4 Tesla K10 GPUs (NVIDIA). The input movie stacks contain 30 sub-frames of 7676×7420 pixels of 8-bit pixel depth. The corrected images were truncated to 3838×3710 pixels in Fourier space before they are saved to disk. For a single stack it took a total of ~62 seconds of which ~42 seconds were spent on computation with the remaining 20 seconds for disk operations. For a batch correction of 10 such stacks, it took ~444 seconds in total. On average the correction of each stack took only 44 seconds. The majority of disk IO time is shadowed by the computational time.

**Cryo-EM data acquisition** Raw micrographs of archaeal 20S proteasome (T. *acidophilum*) and TRPV1 ion channel were acquired in previous studies, as described^6,11^. We also collected a new dataset of archaeal 20S proteasome using Tecnai TF20 electron microscope (FEI Company) equipped with a field emission electron source operating at 200kV acceleration voltage and K2 Summit camera operating in super-resolution mode (Gatan). Frozen hydrated 20S proteasome grids were prepared as previously described. A total of 141 micrographs were collected at a magnification of 29,000X has a physical pixel size of 1.234 Å. The dose rate was set to ~8 e^−^/physical pixel at camera level. Frame exposure time was set to 0.075 second, and total exposure at 10 seconds. corresponding to a total accumulative electron dose of ~65 e^−^/Å^2^ on specimen. The defocus range was set to 0.5 ~ 2.5μm.

**Image processing** Images were subject to motion correction using both MotionCorr and MotionCor2. After motion correction, images were 2x binned by Fourier cropping implemented in both programs. Un-weighted sums were used for CTF determination using CTFFIND4. Weighted sums were used for automated particle picking using RELION and subsequent image processing. The first sub-frame was excluded in both cases. In MotionCor2, we used per-frame dose of 1.36 e^−^/Å^2^ (20S proteasome), 1.30 e^−^/Å^2^ (TRPV1), and 0.58 e^−^/Å^2^ for the new proteasome dataset collected from TF20 microscope for calculating the critical exposure curves for dose-weighting.

Normalized elliptically-averaged Fourier spectrum of un-weighted sum of motion-corrected sub-frames was used to CTF determination using CTFFIND4 ^22^. We used CTFEVAL ^23^ to calculate a cross correlation between the normalized Fourier spectrum and the theoretical CTF, and used 0.5 as a criterion to estimate the resolution of the image. In each dataset, we used 2D class averages calculated from a small number of manually picked particles as references for automatic particle picking using RELION ^24^ 3D reconstructions of each dataset were also calculated and refined following gold-standard refinement procedure implemented in RELION.

For 20S proteasome dataset collected at 300kV, a total of 221,623 particles were autopicked from 570 micrographs corrected with MotionCor2. After reference-free 2D classification, particles from classes that did not depict high-resolution features were excluded, leaving a final set of 187,011 particles. The coordinates of this particle set were used to extract the equivalent particle images from the original MotionCorr corrected micrographs. Initial reference model for 3D reconstruction and refinement was calculated from the atomic structure of archaeal 20S proteasome low-pass filtered to a resolution of 40Å using e2pdb2mrc.py within the EMAN2 package ^25^. The resolutions of final 3D reconstructions with D7 symmetry calculated from the dose-weighted MotionCor2, unweighted MotionCor2, and original MotionCorr particles are 2.58Å, 2.62Å, and 2.73Å respectively. Refined maps were low-pass filtered to the nominal resolution reported and sharpened using the automatic B-factor estimation within the RELION post-processing procedure. Various parameters of the different stages of image processing were kept identical between the un-weighted MotionCor2, dose-weighted MotionCor2, and original MotionCorr particle sets such that differences in resulting maps could be attributed primarily to motion correction and dose weighting.

For the TRPV1 dataset, a total of 236,507 particles were auto-picked from 966 micrographs. All particles were used for reference-based 3D classification implemented in RELION, with C4 symmetry applied. Previously determined TRPV1 density map ^11^ was used as the initial model. A total of 71,825 particles from 3D classes with correct structural features were combined for subsequent 3D refinement. During 3D refinement, we found that applying a soft mask to exclude the density for the cytosolic ankyrin domains during the last five refinement iterations slightly improved the final resolution. Resolutions of final 3D reconstruction calculated from the MotionCor2 dose-weighted, and un-weighted, and original MotionCorr particles are 3.15Å, 3.28Å, and 3.43Å, respectively.

For the 20S proteasome dataset collected with 200kV, image motion was corrected using MotionCor2 only. A total of 36,507 particles were auto-picked and screened by 2D classification from 140 micrographs. Final 3D reconstruction with a D7 symmetry has a resolution of 3.48Å. Furthermore, a 3D reconstruction from a subset of 3,297 particles with defocus between 0.5 – 1.0μm has the resolution of 4.08Å, resolving β-strands within β-sheets. Another 3D reconstruction from a different subset of 3,297 particle with defocus between 2.0 − 2.60μm has the resolution of 4.94Å, where β-strands were not resolved.

***Model Refinement and Visualization*** Atomic model of the 20S proteasome ^6^ was optimized manually in COOT ^26^ and then automatically with phenix.real_space_refine ^27^. All map and model visualization was done in UCSF Chimera.

